# The distribution of neuronal primary *cilia* immunoreactive to melanin-concentrating hormone receptor 1 (MCHR1) in the murine prosencephalon

**DOI:** 10.1101/755967

**Authors:** Giovanne B. Diniz, Daniella S. Battagello, Bianca S. M. Bono, Jozélia G. P. Ferreira, Marianne O. Klein, Livia C. Motta-Teixeira, Jessica C. G. Duarte, Françoise Presse, Jean-Louis Nahon, Antoine Adamantidis, Melissa J. Chee, Luciane V. Sita, Jackson C. Bittencourt

## Abstract

Melanin-concentrating hormone (MCH) is a ubiquitous vertebrate neuropeptide predominantly synthesized by neurons of the diencephalon that can act through two G protein-coupled receptors, called MCHR1 and MCHR2. The expression of *Mchr1* has been investigated in both rats and mice, but its synthesis remains poorly described. After identifying an antibody that detects MCHR1 with high specificity, we employed immunohistochemistry to map the distribution of MCHR1 in the CNS of rats and mice. Multiple neurochemical markers were also employed to characterize some of the neuronal populations that synthesize MCHR1. Our results show that MCHR1 is abundantly found in a sensory subcellular structure called the neuronal primary *cilium*, which has been associated with the detection of free neurochemical agents released to act through volume transmission. Ciliary MCHR1 was found in a wide range of areas, including the olfactory bulb, cortical mantle, *striatum*, hippocampal formation, amygdala, midline thalamic nuclei, periventricular hypothalamic nuclei, and midbrain areas. No differences were observed between male and female mice, and rats and mice diverged in two key areas: the caudate-putamen nucleus and the subgranular zone of the dentate gyrus. Ciliary MCHR1 was found in close association to several neurochemical markers, including tyrosine hydroxylase, calretinin, kisspeptin, estrogen receptor, oxytocin, vasopressin, and corticotropin-releasing factor. Given the role of neuronal primary *cilia* in sensing free neurochemical messengers in the extracellular fluid, the widespread distribution of ciliary MCHR1, and the diverse neurochemical populations who synthesize MCHR1, our data indicates that volume transmission may play a prominent role in the normal function of the MCH system.

## 1. Introduction

Melanin-concentrating hormone (MCH) is a ubiquitous vertebrate neuropeptide, synthesized predominantly by neurons of the diencephalon and, in particular, of the lateral hypothalamic area (Bittencourt & Diniz 2018; Bittencourt *et al.* 1992). The MCH neuropeptidergic system has been involved in several physiological roles, including the integration and promotion of motivated behaviors (Diniz & Bittencourt 2017), modulation of energy expenditure (Qu *et al.* 1996; Chee *et al.* 2019), sleep (Ferreira *et al.* 2017a; Gao 2018), ventricular homeostasis (Conductier *et al.* 2013b; Conductier *et al.* 2013a), and autonomic function modulation (Brown *et al.* 2007; Messina & Overton 2007). It has also been implicated in reproductive physiology (Naufahu *et al.* 2013), including the release of luteinizing hormone (Chiocchio *et al.* 2001; Murray *et al.* 2000b), the promotion of sexual behavior (Gonzalez *et al.* 1996), and parental behavior (Alachkar *et al.* 2016; Benedetto *et al.* 2014).

The mammalian MCH peptidergic system is composed by three peptides produced from a single precursor, encoded by the *Pmch* gene. In addition to MCH, neuropeptide E-I (NEI) and neuropeptide G-E (NGE) are cleaved by prohormone convertases and may have biological activities that are independent of MCH (Bittencourt & Celis 2008). This system is also composed by two receptors that bind with specificity to MCH, called MCH receptor 1 (MCHR1) and receptor 2 (MCHR2) (reviewed in Presse *et al.* (2014) and Bittencourt and Diniz (2018)). While MCHR1 is present in a wide range of vertebrates, MCHR2 has been lost after the divergence of the *Glires* clade, what resulted in the loss of a working MCHR2 receptor in rabbits, guinea pigs, hamsters, rats and mice (Tan *et al.* 2002). As a result, the amount of information amassed about MCHR1 is significantly higher than that about MCHR2.

Despite the focus on MCHR1, there are a few important aspects of this receptor that are not well understood. After the discovery of MCH as the ligand of MCHR1 (Bächner *et al.* 1999; Chambers *et al.* 1999; Saito *et al.* 1999; Shimomura *et al.* 1999), the distributions of *Mchr1* gene expression and MCHR1 immunoreactivity were described in rats (Saito *et al.* 2001; Hervieu *et al.* 2000). In both cases, only male specimens were used, and both works describe widespread expression/synthesis of MCHR1. The next wave of works then focused on mice, through the use of gene reporters (Chee *et al.* 2013; Engle *et al.* 2018). Despite an exhaustive investigation and detailed description of the gene reporter distribution, the experimental model of gene reporters cannot inform the subcellular localization of the protein product that results from the investigated gene. It is unclear, therefore, what is the precise subcellular localization of MCHR1 *in vivo.*

The first hint that the subcellular localization of MCHR1 may be important for its function came in the work of Berbari *et al.* (2008), who described MCHR1 immunoreactivity in the primary *cilia* of cultured cells, and in a few areas of the central nervous system. Primary *cilia* are non-motile structures, found within the CNS exclusively in neurons, and first described in humans (Mandl & Megele 1989). The main role of primary *cilia* is chemical sensing and signal transducing (Pazour & Witman 2003; Berbari *et al.* 2009), as they are covered in receptors (Händel *et al.* 1999; Brailov *et al.* 2000; Loktev & Jackson 2013; Koemeter-Cox *et al.* 2014; Siljee *et al.* 2018), and participate in several signaling pathways (Breunig *et al.* 2008; Rohatgi *et al.* 2007; Corbit *et al.* 2005). Neuronal primary *cilia* may be an integral part of volume transmission (VT), a neuronal method of communication that involves the use of neurochemical messengers outside the synapse (Agnati *et al.* 1986; Agnati *et al.* 1995; Agnati *et al.* 2010), including the release of neuroactive substances in the extracellular space (ECS) and the cerebrospinal fluid (CSF) (Fuxe *et al.* 2007; Agnati & Fuxe 2014).

Mapping the areas where MCHR1 is associated to primary *cilia* in the CNS may help us understand its mechanisms of action. It has recently been demonstrated that MCH acts in a VT paradigm to exert a modulatory role in feeding (Noble *et al.* 2018; Jiang & Brüning 2018). Areas linked to feeding behavior where MCHR1 is ciliary are the likely target for this effect. Furthermore, areas unrelated to feeding behavior that contain ciliary MCHR1 may reveal clues about other actions exerted by MCH that depend on VT. With that in mind, we mapped the presence of MCHR1 immunoreactivity in *cilia* in the prosencephalon of adult rats and mice. Since MCH has been implicated in sexual regulation (Gonzalez *et al.* 1996; Murray *et al.* 2000b; Murray *et al.* 2006; Wu *et al.* 2009), and female models have been poorly explored in the literature concerning MCH, we included females in all four estrous cycle stages to evaluate possible changes in the receptor distribution linked to reproductive status. The results found in mice were then compared to rats to inform us about possible interspecies differences in the distribution of MCHR1.

## 2. Materials and Methods

### 2.1. Animals

Adult male and female virgin C57BL/6 mice (n = 25) were bred and raised in the animal facility of the Department of Anatomy, Institute of Biomedical Sciences of the University of São Paulo. At the beginning of the experiments, animals were approximately three months old and weighed between 20 and 30g. Females were housed five per cage, while males were housed individually after they reached sexual maturity, in a room with 12/12h light/dark cycle (lights on at 6:00 AM), controlled temperature (22±2°C), and *ad libitum* access to water and standard chow. All experiments were carried out in accordance with the Guidelines for the Care and Use of Mammals in Neuroscience and Behavioral Research established by the Brazilian National Research Council (CONCEA, 2016), as well as with those established by the Ethics Committee on the Animal Use of the Institute of Biomedical Sciences (Protocol CEUA #096/2014). No effort was spared to minimize the number of animals employed in this study and any incidental suffering caused to them. To evaluate the possibility that interspecies differences exist in the distribution of MCHR1, adult male Sprague-Dawley rats (n = 5) were bred and raised as described above. To test antibody specificity, we employed *Mchr1^-/-^* mice (Adamantidis *et al.* 2005), as well as their heterozygous and WT littermates. These animals were bred and raised in the animal facility of the University Hospital, Inselspital, in Bern, Switzerland in conditions similar as above.

To determine the day of euthanasia, females had their estrous cycle checked daily at 10:00 AM by the method of vaginal cytology (Byers *et al.* 2012). To that end, 50μl of saline were inserted in the vaginal channel with a pipette and gel-loading tips. Care was taken so the tip did not touch the vagina to avoid unintended stimulation and pseudopregnancy. The liquid was then collected, spread on glass slides, and examined under the light microscope (Nikon Corporation; Minato, Tokyo, Japan). Estrous cycle determination was based on the relative densities of epithelial, cornified, and leucocyte cell populations (Byers *et al.* 2012).

### 2.2. Tissue collection

At the day of euthanasia, animals were anesthetized with an excess of Xylazine, Ketamine and Acepromazine. Once verified the loss of reflexes, animals were thoracotomized and transcardially perfused. For mice, approximately 20ml of cold 0.9% saline solution were perfused to clean the vascular bed, followed by 240ml of 4% formaldehyde solution in borate buffer (pH 9.5). In rats, the volume was 100ml of cold 0.9% saline and 750 ml of 4% formaldehyde solution in borate buffer (pH 9.5). After perfusion, the heads were separated from the body, the cranium opened, and the brain removed and post-fixed in the same fixative solution with 20% sucrose for four hours. At the end of the postfixation time, brains were transferred to 0.02M potassium-phosphate buffered saline (KPBS, pH 7.4) with 20% sucrose at 4°C until slicing. Slicing of the brains was performed in the frontal plane in a sliding microtome. One-in-five series of 20μm slices were obtained from mice and one-in-five series of 40μm were obtained from rats. Slices were kept in antifreeze solution at −30°C until immunohistochemistry.

### 2.3. Immunohistochemistry

#### 2.3.1. Antibody selection

To identify an antibody that selectively labels MCHR1, we tested six brands of commercially-available antibodies. The details of each antibody are listed in Table 1. Each antibody was used for immunoperoxidase staining in six different dilutions (1:1,000, 1:3,000, 1:10,000, 1:30,000, 1:100,000 and 1:300,000), as described by Hoffman *et al.* (2016) for antibody titration. In some cases, where no signal whatsoever could be detected, we tested more concentrated titers (not shown). The concentration of antibody found to be optimal for immunoperoxidase staining was then used for immunofluorescence procedures. Primary and secondary antibody suppression tests were performed for all antibodies (not shown). The antibody that resulted in the best labeling was then further tested on tissue from knockout animals, negative control test described as the gold standard of antibody validation by Saper and Sawchenko (2003).

**Table 1.** Primary antibodies employed in this work

| Antibody | Manufacturer | Man. ID | RRID | Antibody | Manufacturer | Man. ID | RRID |
| --- | --- | --- | --- | --- | --- | --- | --- |
| Rabbit anti melanin-concentrating hormone receptor 1 (anti-hMCHR1) | ThermoFisher Scientific | PA5-24182 | AB_2541682 | Rabbit gonadotropin-releasing hormone (anti-GnRH) | ImmunoStar | 20075 | AB_572248 |
| Rabbit anti melanin-concentrating hormone receptor 1 (anti-hMCHR1) | Abcam | Ab97509 | AB_10680290 | Rabbit anti kisspeptin (anti-KISS) | Millipore (Chemicon) | AB9754 | AB_2296529 |
| Rabbit anti melanin-concentrating hormone receptor 1 (anti-hMCHR1) | SantaCruz Biotechnology | sc-25667 | AB_2143948 | Rabbit anti estrogen receptor $\alpha$ (anti-ERα) | Millipore (Upstate) | 06-935 | AB_310305 |
| Rabbit anti melanin-concentrating hormone receptor 1 (anti-hMCHR1) | SantaCruz Biotechnology | Sc-5534 | AB_2143957 | Rabbit anti vasopressin (anti-AVP) | ImmunoStar (DiaSorin) | 20069 | AB_572219 |
| Rabbit anti melanin-concentrating hormone receptor 1 (anti-hMCHR1) | Sigma-Aldrich/Atlas | HPA004149 | AB_1079363 | Guinea-pig anti oxytocin (anti-OT) | Peninsula | T-5021.0050 | AB_518526 |
| Rabbit anti melanin-concentrating hormone receptor 1 (anti-rMCHR1) | Alomone Labs | AMR-041 | AB_11218957 | Rabbit anti corticotropin-releasing factor (anti-CRF) | Peptide Biology Laboratory, The Salk Institute | rC70 | AB_2650437 |
| Rabbit anti adenylate cyclase III (anti-AC3) | ThermoFisher Scientific | PA5-35382 | AB_2552692 | Sheep anti α-melanocyte-stimulating hormone (anti-α MSH) | Millipore (Chemicon) | AB5087 | AB_91683 |
| Rabbit anti melanin-concentrating hormone (anti-MCH) | Peptide Biology Laboratory, The Salk Institute | PBL #234 | AB_2650444 | Rabbit anti cocaine- and amphetamine-regulated transcript (anti-CART) | Peptide Biology Laboratory, The Salk Institute | 6838 | AB_2650446 |
| Mouse anti tyrosine hydroxylase (anti-TH) | ImmunoStar | #22941 | AB_572268 | Mouse anti neuron-specific nuclear protein (anti-NeuN) | Millipore (Chemicon) | MAB377 | AB_2298772 |
| Mouse anti calbindin D-28k | Swant | 300 | AB_10000347 | Mouse anti glial fibrillary acidic protein (anti-GFAP) | Sigma-Aldrich | G3893 | AB_477010 |
| Goat anti (rat, mouse, human) calretinin | Swant | CG1 | AB_10000342 | Mouse anti doublecortin (anti-DCX) | Santa Cruz | Sc-8066 | AB_2088494 |

#### 2.3.2. Immunoperoxidase

For immunoperoxidase staining, slices were extensively rinsed in KPBS for antifreeze solution removal and incubated with a 0.3% solution of hydrogen peroxidase in KPBS for 15 minutes. Slices were then rinsed again in KPBS and incubated with primary antibody in a KPBS solution of 3% normal goat serum (Vector Laboratories; Burlingame, CA, USA; AB_2336615) and 0.3% Triton X-100 for membrane permeabilization. In the following day, slices were rinsed in KPBS and incubated with biotinylated goat anti-rabbit IgG (1:800, Vector; AB_2313606) for one hour at room temperature. After new rinses, slices were incubated with an avidin-biotin-horseradish peroxidase solution (1:333, Vector; AB_2336819) for one hour. For chromogen deposition, slices were transferred to a solution of 3,3’-diaminobenzidine tetrahydrochloride (DAB, 0.02% – Sigma Chemical; St. Louis, MO, USA) with 0.003% hydrogen peroxide and 2.5% nickel ammonium sulfate diluted in 0.2M sodium acetate buffer (pH 6.5). Reactions were stopped by transferring slices to clean sodium acetate buffer and then rinsing them with KPBS. Sections were then mounted on gelatin-coated glass slides in rostrocaudal order, dehydrated, defatted and coverslipped with a hydrophobic mounting medium (DPX; Sigma).

#### 2.3.3. Immunofluorescence

For immunofluorescence staining of MCHR1, a protocol similar to that described by Hoffman et al (2016) was employed. Briefly, slices were rinsed in KPBS and incubated with a 0.3% solution of hydrogen peroxidase in KPBS for 20 minutes. After new rinses, slices were incubated with primary antibody in a KPBS solution of 3% normal donkey serum (Jackson ImmunoResearch; West Grove, PA, USA; AB_2337258) and 0.3% Triton X-100 for membrane permeabilization. In the following day, slices were incubated with biotinylated donkey anti-rabbit IgG antibody (1:5000, Jackson ImmunoResearch; AB_2340593) for one hour at room temperature and then incubated with avidin-biotin-horseradish peroxidase solution (1:833, Vector; AB_2336819) for 30 minutes at room temperature. Slices were then washed and incubated with a 0.5% biotinylated (EZ-Link sulfo-NHS-LC-biotin; ThermoFisher Scientific, Waltham, MA, USA) tyramine (Sigma-Aldrich; Sigma Chemical; St. Louis, MO, USA) and hydrogen peroxide (0.005%) solution for 20 minutes, and then incubated with AlexaFluor 594-conjugated streptavidin (1:200, Invitrogen; Carlsbad, CA, USA). Slices were then rinsed in KPBS, incubated with DAPI (1:10’000, Invitrogen) for 10 minutes and mounted in glass slides or used in following labeling experiments.

To label other markers, MCHR1-immunolabeled slices were rinsed with KPBS and incubated with primary antibodies for different neurochemical markers (Table 1) in KPBS containing 3% normal donkey serum (Jackson ImmunoResearch; AB_2337258) and 0.3% Triton X-100. In the following day, slices were rinsed and incubated with AlexaFluor 488-conjugated donkey anti-IgG antibodies (1:200, Invitrogen; AB_2556546, AB_2534082, AB_2556542, AB_2534102) for two hours at room temperature. After final rinses with KPBS, slices were mounted in glass slides. After a brief period of air drying, the slides were then covered with antifade mounting medium (Invitrogen) and coverslipped.

### 2.4. Imaging and Data Analysis

Immunoperoxidase-labeled slices were examined with brightfield microscopy in a Nikon 80i microscope (Nikon) coupled to a digital camera (CX3000 – MBF Bioscience Co.; Williston, VT, USA) and operating under the software Microlucida 3.03 (MBF Bioscience). Immunofluorescence-labeled slices were examined with widefield and confocal fluorescence microscopy in an AxioImager Z2 motorized upright microscope (Carl Zeiss; Wetzlar, Germany). Widefield illumination was obtained with an HXP 120V illuminator (Carl Zeiss) and photomicrographs were produced with an AxioCam 506c (Carl Zeiss). Each fluorophore was imaged separately in emulated monochromatic mode in separate channels and then merged. Confocal microscopy was performed using a laser unit as light source and the LSM800 scan unit with two metal halide detectors (Carl Zeiss). All images had adjustments in their brightness, contrast, and sharpness applied to every pixel in the picture, and the changes did not alter the information illustrated in the figures. The plates were assembled, and off-tissue background/methodological artifacts were cleared by means of Adobe Photoshop CC 2017.0.1 software (Adobe Systems Inc; San Jose, CA, USA; SCR_014199).

Mapping of MCHR1 immunoreactivity was performed by comparing the cytoarchitectonic characteristics of each slice, as evidenced by DAPI staining (when applicable), to the atlas of Paxinos and Franklin (2012) and Paxinos and Watson (2006). In some cases, additional information was obtained using neurochemical markers to delineate specific areas. The comparison of relative densities between male and female and between different estrous cycle stages was performed using a semi-quantitative method. After analysis of random slices, a scale of *cilia* density was developed. An investigator blind to the experimental condition of the animals then used the scale to rate the presence of immunoreactive *cilia* in each area examined with a number ranging from 0-8, with 0 corresponding to complete absence of immunoreactivity and 8 corresponding to the highest density found. The same procedure was performed for all animals and the average density of *cilia* was obtained for each area and each sexual status. To facilitate the presentation of these results, the number system was then converted to a system of crosses with less discrimination but better noise suppression.

## 3. Results

### 3.1. The antibody used targets ciliary MCHR1 with high specificity

From the six antibodies tested for MCHR1 detection (Table 1), only one resulted in consistent labeling at the tested concentrations: antibody PA5-24182 from Invitrogen. Antibody Santa Cruz C-17 (Table 1) also resulted in staining, but the lower signal/background *ratio* and the discontinuation of this antibody by its manufacturer led us to use antibody PA5-24182 in the rest of the experiments. Because antibody PA5-24182 has never been used in the literature, we proceeded to verify its specificity by employing standard omission tests and using tissue from *Mchr1^-/-^* and *Mchr1^+/-^* mice (Saper & Sawchenko 2003). As expected, antibody and fluorophore omission tests were successful, with no staining observed in the antibody suppression cases (Fig. 1 A’-A”). Using the antibody on tissue from *Mchr1^+^* animals resulted in the regular pattern of staining observed during titrations (Fig. 1 A, B). No staining could be observed on slices from *Mchr1^-/-^* animals (Fig. 1 B”). Interestingly, *Mchr1^+/-^* animals displayed an intermediate pattern of staining, with scattered labeling found in areas otherwise heavily labeled in *Mchr1^+^* animals (Fig. 1 B’). These tests show that the used MCHR1 antibody labels its target with high specificity.

**Figure 1 –.**
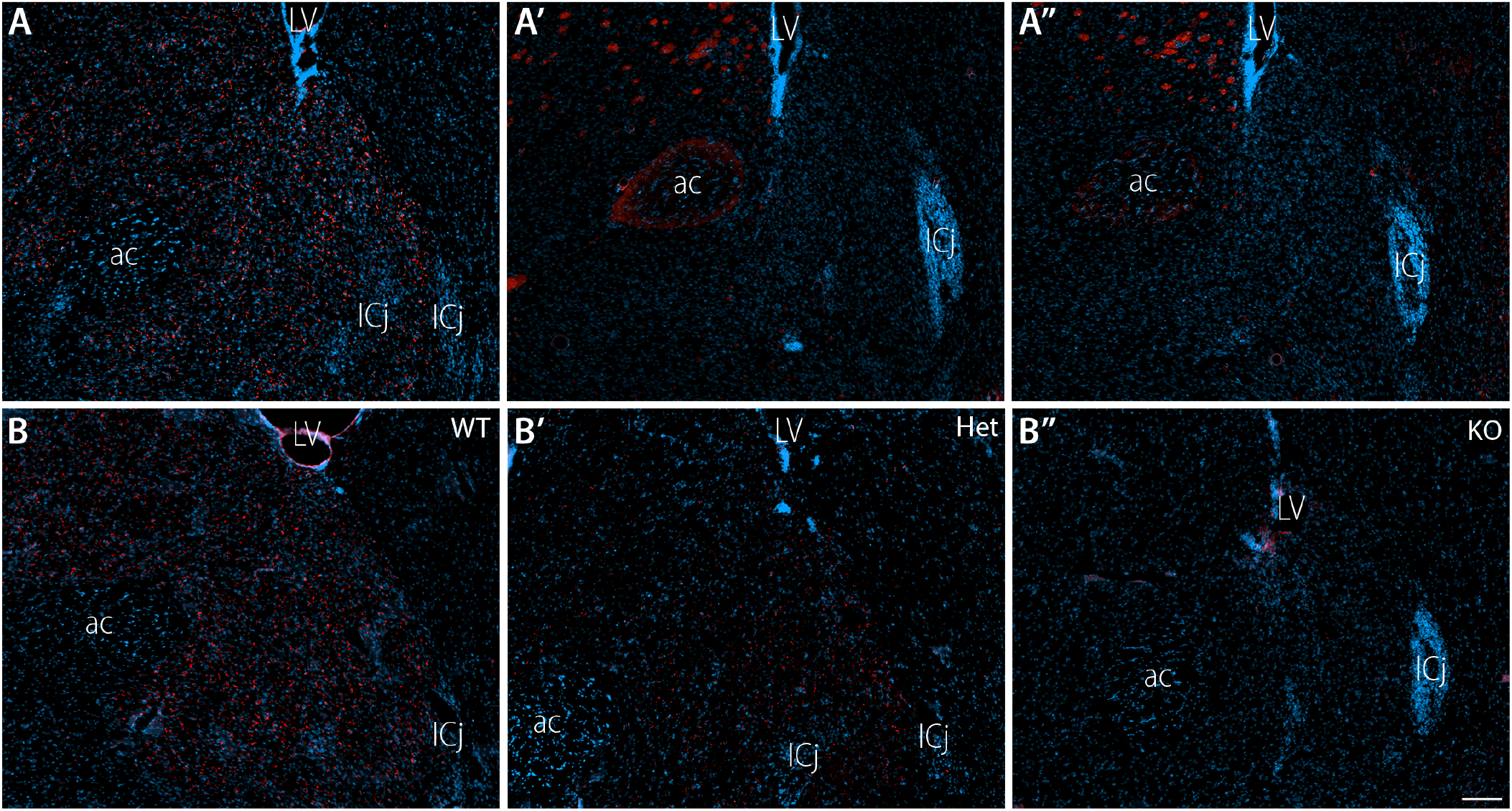
Specificity controls for the MCHR1 antibody employed in this work. Widefield fluorescence photomicrographs of frontal brain slices obtained from male mice with different genotypes submitted to immunohistochemistry with or without antibodies and counterstained with DAPI (blue). A) Reaction in the presence of primary and secondary antibodies results in ample labeling of the *nucleus accumbens*; A’) Omitting the primary antibody results in no labeling of *cilia* in the *nucleus accumbens.* Note that red fluorescence on the anterior commissure and matrix of the caudate-putamen are caused by overexposure of the red channel, and do not represent specific labeling; A’’) As is the case with the primary antibody, omitting the secondary biotinylated antibody also resulted in abolished labeling; B) The use of antibody PA5-24182 results in ample labeling of *cilia* in WT C57BL/6 mice; B’) In heterozygous mice, a clear reduction in the number of labeled *cilia* is observed; B’’) In homozygous Mchr1^-/-^ mice, no labeling can be observed. Abbreviations: ac – anterior commissure; ICj – islands of Calleja; LV – lateral ventricle. Scale bar: 100μm.

To confirm the labeling observed using the PA5-24182 was ciliary, we performed double immunohistochemistry in series of some animals using an anti-adenylate cyclase III (AC3) antibody. Adenylate cyclase III has been shown to be a marker of neuronal primary *cilia* (Bishop *et al.* 2007). As expected, we found widespread staining for AC3, agreeing to the distribution described by Bishop et al (2007). In several areas, colocalization between MCHR1 and AC3 was almost complete, such as in the pyramidal layer of the hippocampus proper (Fig. 2 A), while in others there was no signal of MCHR1 colocalizing with the AC3-positive labeling, such as in the granular layer of the dentate gyrus (Fig. 2 B). Upon closer inspection, it was observed that MCHR1 labeling seldom covers the whole *cilia*, in most cases forming a patched pattern upon the ciliary surface (Fig. 2 A’). As an additional control, we performed simultaneous MCHR1 and NeuN (neuronal marker) or GFAP (glial marker) labeling. Since neuronal primary *cilia* are not found in glial cells, we expected no clear pattern of co-distribution between MCHR1 and GFAP. As expected, no clear pattern was observed, while MCHR1 was found codistributed with NeuN in several areas (Supplementary Material 01).

**Figure 2 –.**
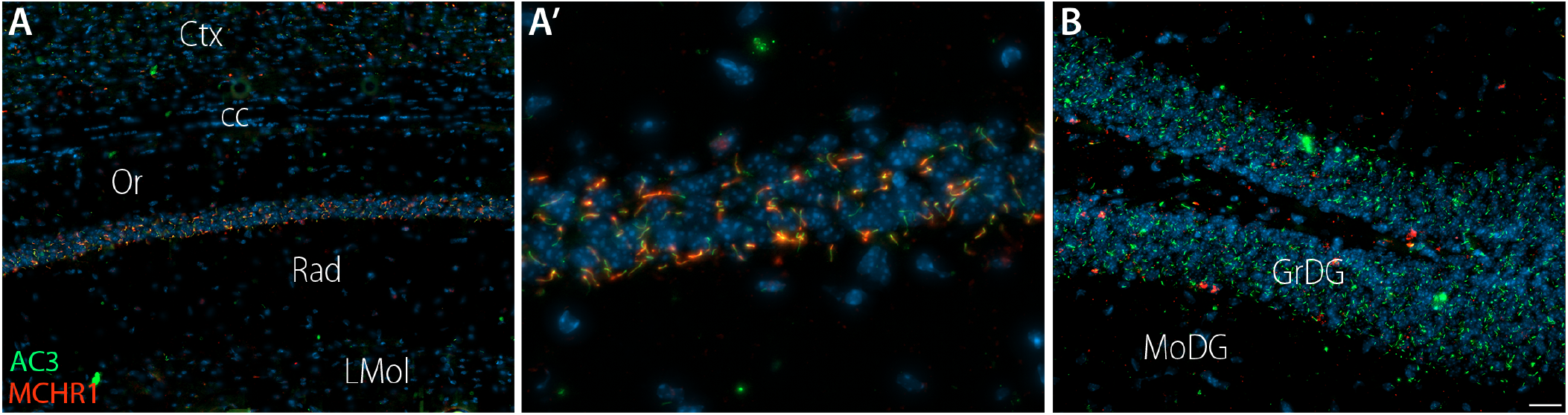
The subcellular localization of MCHR1. Widefield fluorescence photomicrographs of frontal male mouse brain slices submitted to immunohistochemistry for adenylate cyclase III (green) and MCHR1 (red) and counterstained with DAPI (blue). A) In the pyramidal layer of the hippocampus, there is a large degree of colocalization between MCHR1 and AC3, as revealed by the merged yellow-orange labeling; A’) Higher magnification of A. Immunoreactivity to MCHR1 is found in restricted areas of the AC3-positive *cilia,* varying in terms of coverage of the ciliary surface; B) Other areas, such as the granular layer of the dentate gyrus, are rich in AC3-ir *cilia,* but lack any MCHR1. Abbreviations: cc – *corpus callosum*; Ctx – cortex; GrDG – granular layer of the dentate gyrus; LMol – *stratum lacunosum moleculare* of the hippocampus; MoDG – molecular layer of the dentate gyrus; Or – *stratum oriens* of the hippocampus; Rad – *stratum radiatum* of the hippocampus. Scale bar: A, B = 100 μm; A’ = 50μm.

### 3.2. MCHR1 is widely distributed within the murine prosencephalon

To map the distribution of MCHR1 in the mouse prosencephalon, we developed a grading scale that went from 0 (complete absence of staining) to 8 (maximum density of weight) (Fig. 3). Each area identified using the mouse brain atlas by Paxinos and Franklin (2012) from each animal was then scored based on that grading scale by a single experimenter, who was blind to the group of each animal. Upon comparison, there were no consistent differences between male and female and the different stages of the estrous cycle, and therefore only a stereotypical distribution contemplating all groups will be presented in this paper (Table 2). The distribution of MCHR1 in rats and mice was also highly similar, and therefore the mouse will be used to elaborate the description of the results, and differences between the two species will be highlighted at the end of this section. To simplify the description of the results, the numerical system implemented in the first analysis was simplified to a cross system, where a dash represents no staining and four crosses represent maximum staining (Fig. 3). Highlights of the morphological distribution of MCHR1-ir *cilia* in adult mice will be provided in this section, and some areas are illustrated in Fig. 4. For an exhaustive description, we ask the reader to refer to Table 2.

**Figure 3 –.**
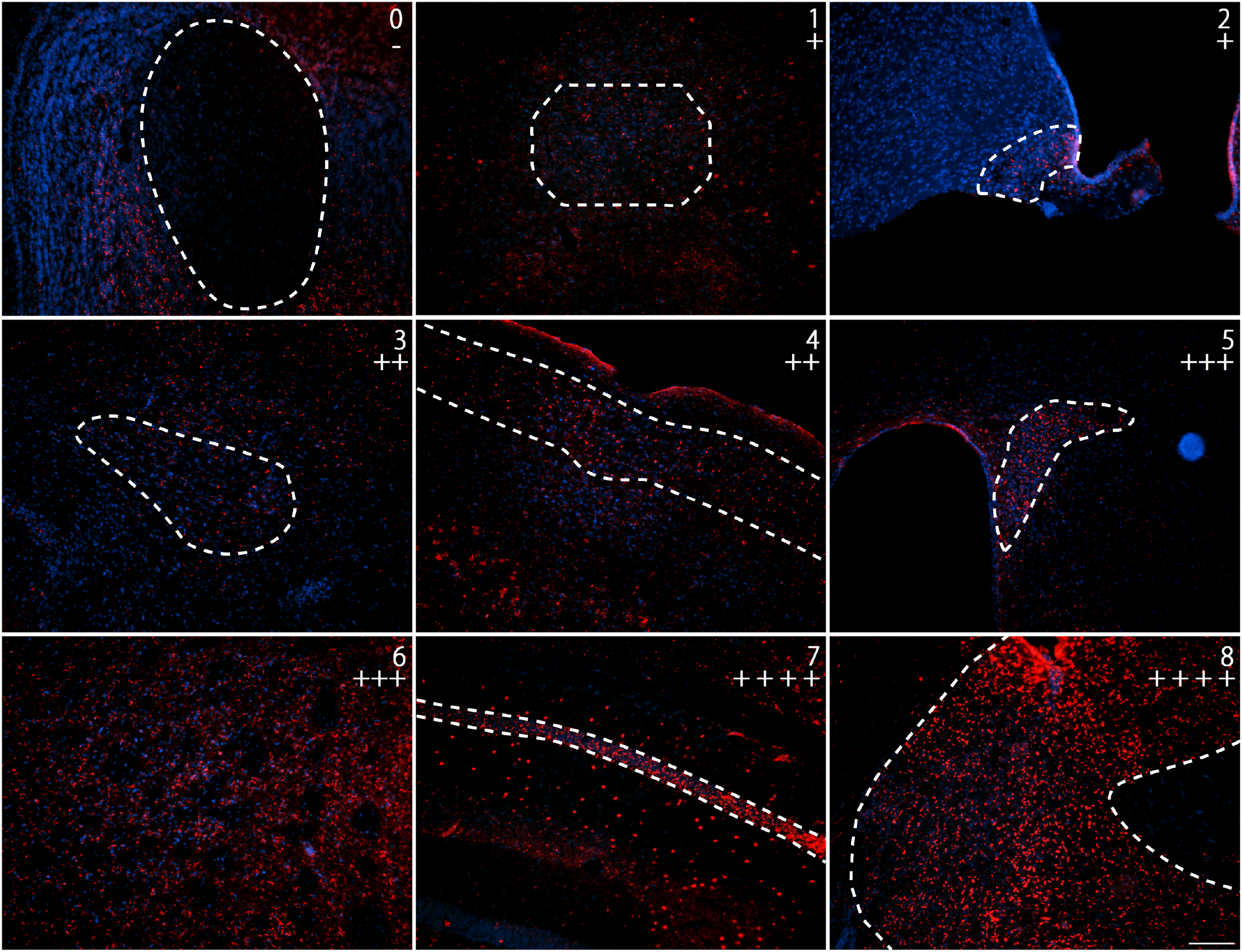
Scale of MCHR1 immunoreactivity density. Widefield fluorescence photomicrographs of frontal mice brain slices submitted to immunohistochemistry for MCHR1 (red) and counterstained with DAPI (blue). The areas used to elaborate this scale are: accessory olfactory bulb (0 | −), central medial thalamic nucleus (1 | −/+), arcuate nucleus (2 | +), basolateral nucleus of the amygdala (3 | ++), cortical layers 2 and 3 (4 | ++), paraventricular nucleus (5 | +++), caudate-putamen (6 | +++), pyramidal layer of the hippocampus proper (7 | ++++), and shell of the *nucleus accumbens* (8 | ++++). Scale bar: 100μm.

**Figure 4 –.**
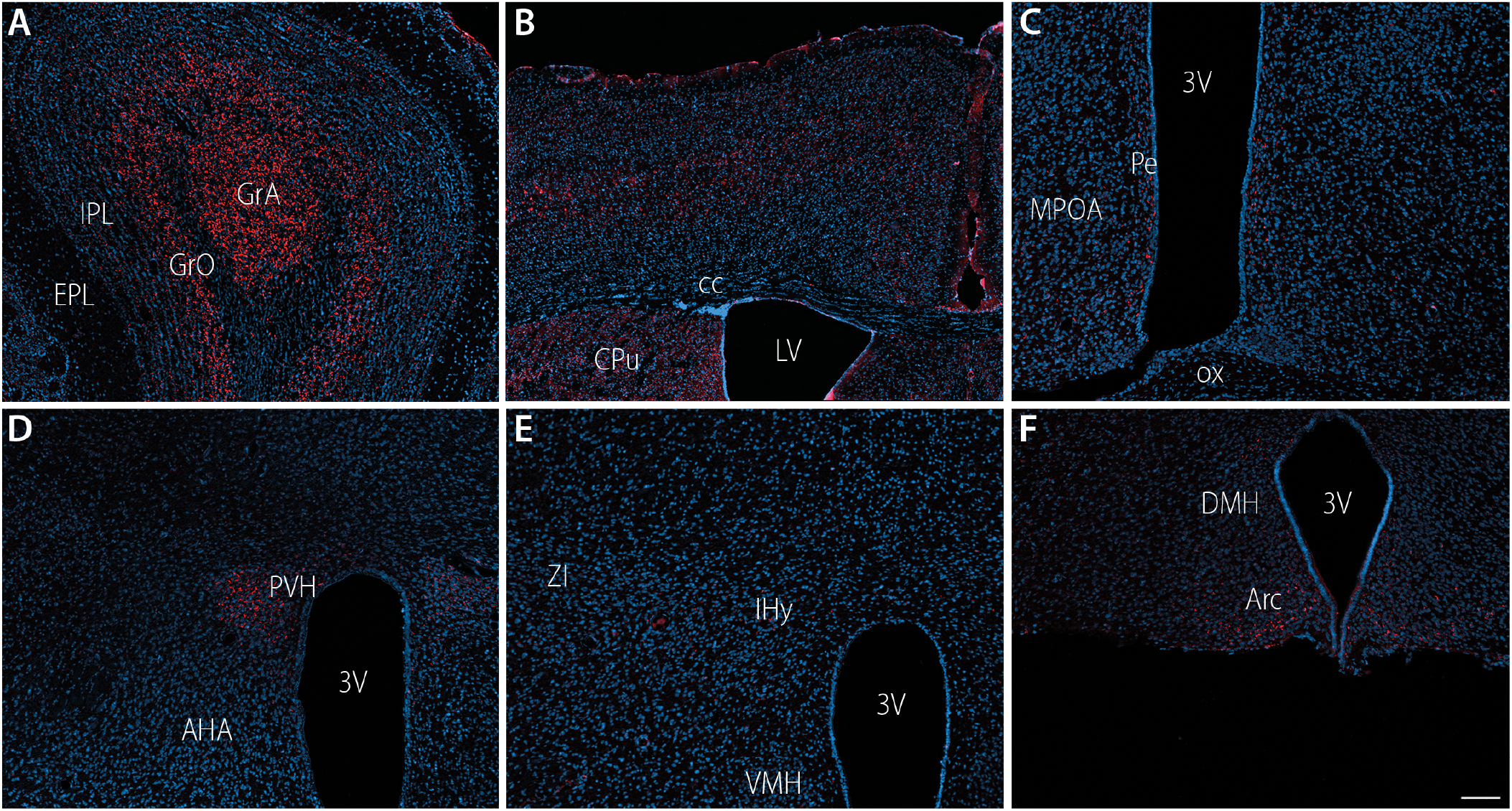
MCHR1 immunoreactivity in discrete prosencephalic areas. Widefield fluorescence photomicrographs of frontal mice brain slices submitted to immunohistochemistry for MCHR1 (red) and counterstained with DAPI (blue). A) The olfactory bulb contains one of the highest densities of ciliary MCHR1 in the mouse prosencephalon; B) Moderate to high densities of ciliary MCHR1 are found in layers 2, 3 and 5 of several cortical areas. A high density of labeling is observed in the caudate-putamen nucleus at this same level; C) A moderate density of *cilia* is observed in the preoptic periventricular nucleus of the hypothalamus. Some labeled *cilia* are also observed in the medial preoptic area; D) the paraventricular hypothalamic nucleus harbors one of the highest numbers of MCHR1-labeled *cilia* in the hypothalamus; E) Although only a few neurons contain neurons with MCHR1-harboring *cilia* in the incerto-hypothalamic area, this area can be clearly identified due to the lack of labeled *cilia* around it; F) The arcuate nucleus also contains a large number of immunolabeled *cilia* when compared to other hypothalamic nuclei, in particular in its ventromedial portion. Abbreviations: 3V – third ventricle; AHA – anterior hypothalamic area; Arc – arcuate nucleus; cc – *corpus callosum*; CPu – caudate-putamen; EPL – external plexiform layer of the olfactory bulb; GrA – granular layer of the accessory olfactory bulb; GrO – granular layer of the olfactory bulb; IPL – internal plexiform layer of the olfactory bulb; IHy – incerto-hypothalamic area; LV – lateral ventricle; MPOA – medial preoptic area; Pe – periventricular preoptic nucleus; PVH – paraventricular hypothalamic nucleus; VMH – ventromedial hypothalamic nucleus; ZI – *zona incerta.* Scale bar: 100μm.

**Table 2.**
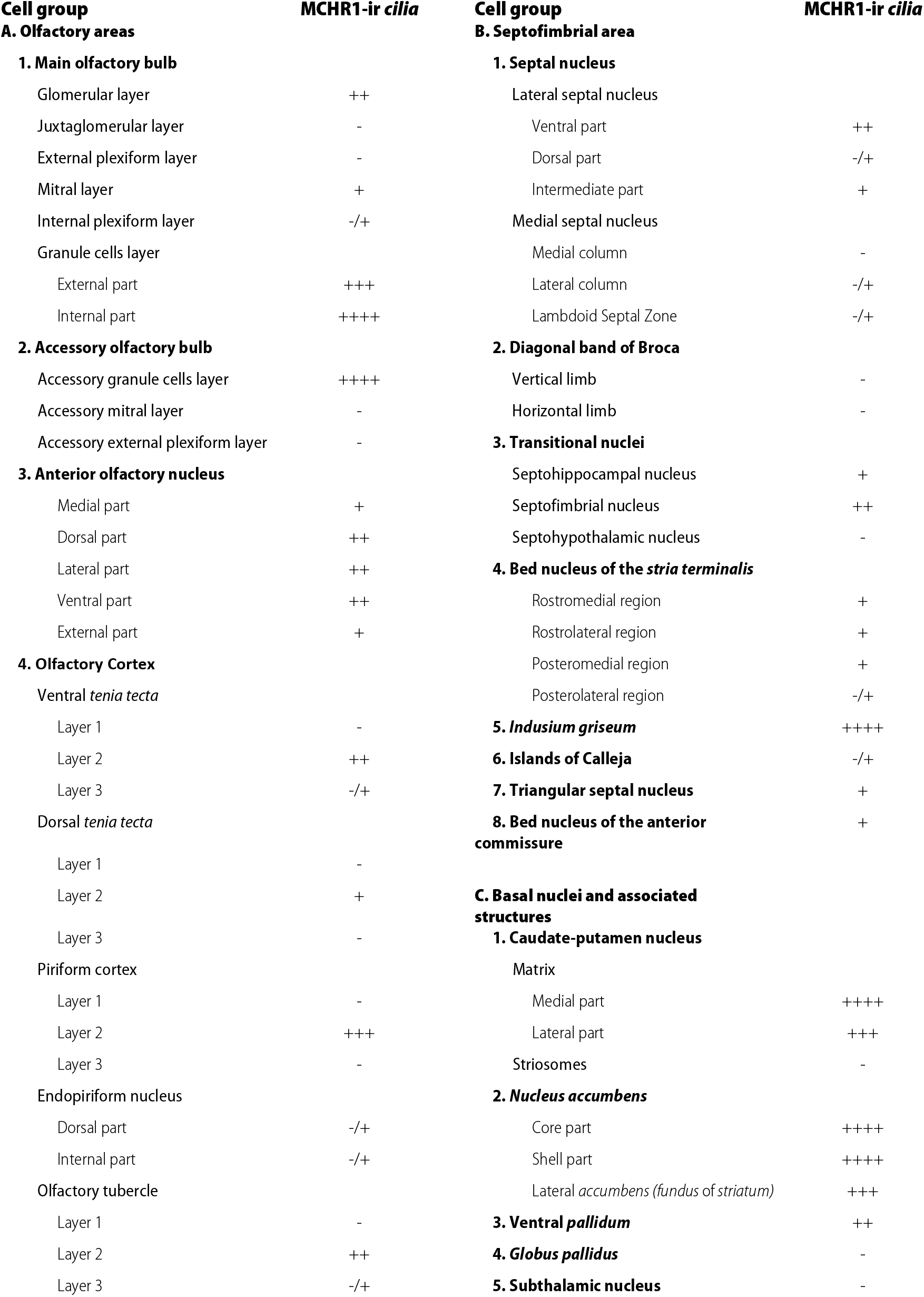
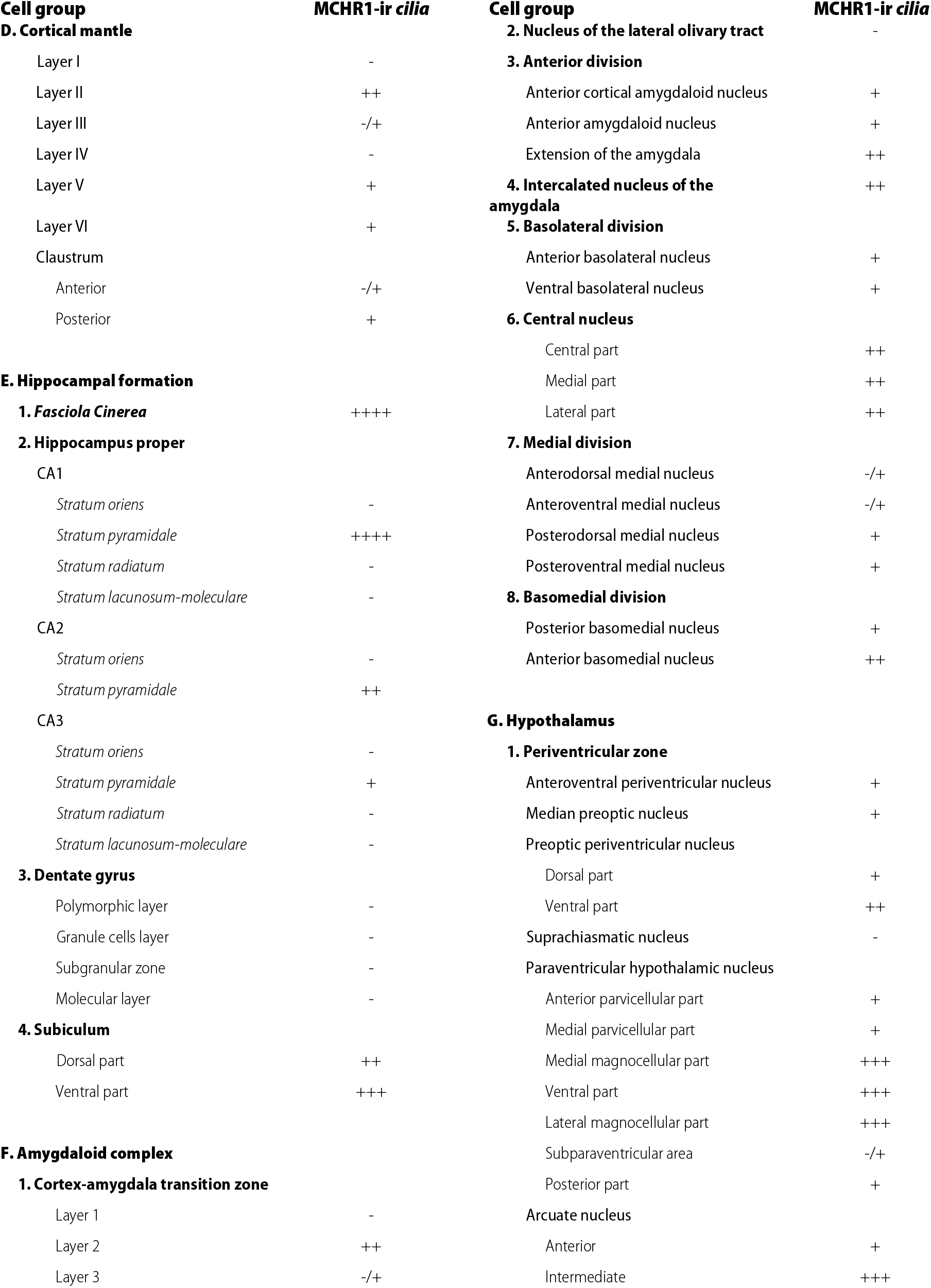
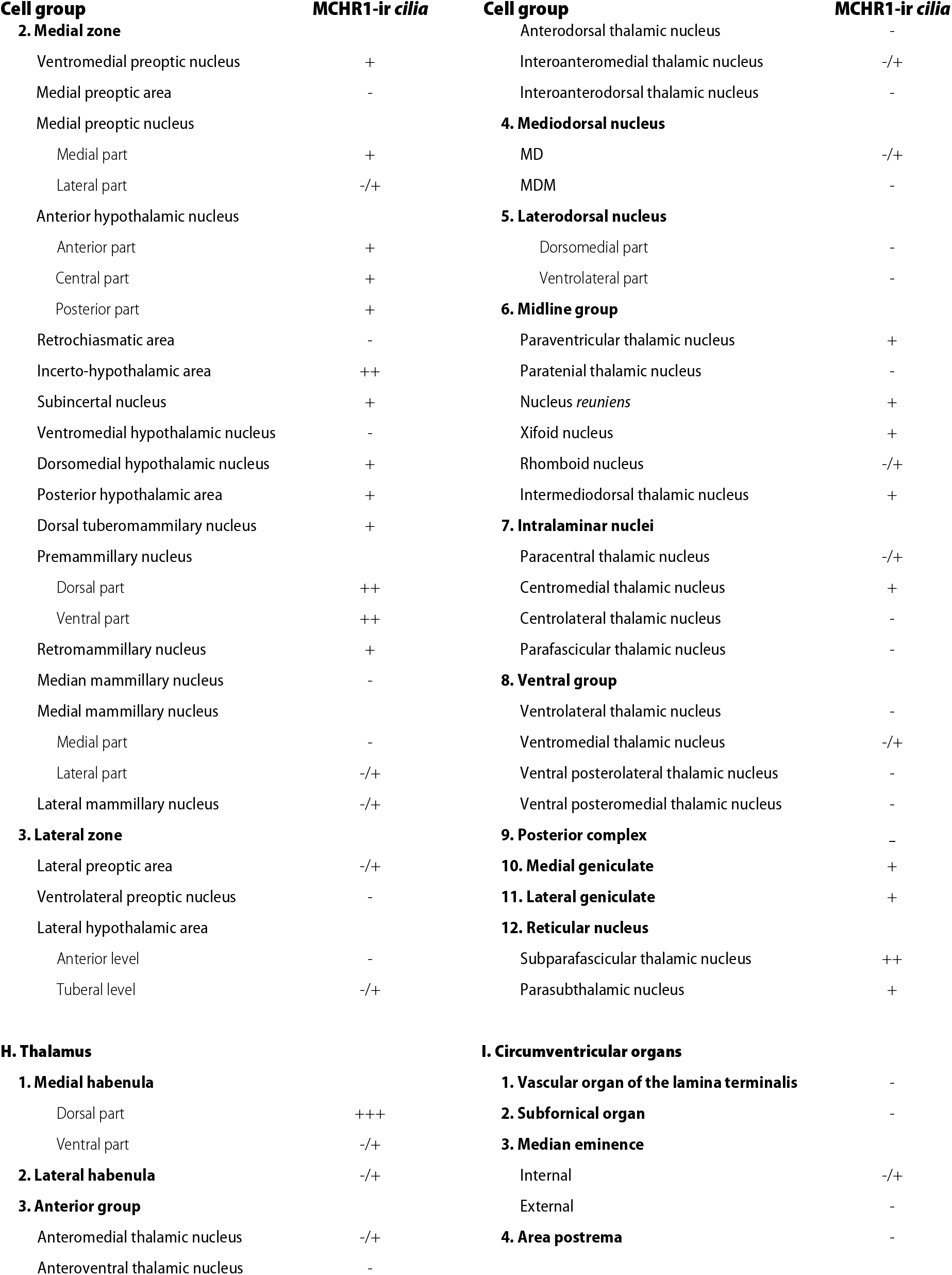
Relative density of MCHR1-ir *cilia* in the mouse CNS

Labeling could be abundantly observed in olfactory areas, such as the olfactory bulb, including the glomerular and granular layers, internal plexiform layer and mitral cell layer; layer 2 of the piriform cortex; dorsal and ventral *taenia tecta;* dorsal and intermediate endopiriform nucleus; and olfactory tubercle. Albeit in a smaller density, we also observed labeling in the medial aspect of the anterior olfactory nucleus. Other subcortical areas were also rich in labeling. The region with the densest immunolabeling for MCHR1 was the shell of the *nucleus accumbens,* with a moderate density found in the core subdivision of this nucleus. The neighboring *ventral pallidum* also displayed high levels of MCHR1 immunoreactivity. In the caudate-putamen nucleus, a mediolateral gradient was observed in the matrix, with a high density found medially, close to the lateral ventricle walls. No staining was observed in the striasomes. Likewise, no staining was observed in the *globus pallidus*. In the adjoining septal nuclei, only sparse immunoreactivity was seen in the ventral part of the lateral septal nucleus, with no labeling detected in other subdivisions of the lateral septal nucleus or in the medial septal nucleus.

In the amygdaloid complex, scattered MCHR1 immunoreactivity was observed in the basolateral, basomedial, medial and central nuclei. The hippocampal formation displayed a very characteristic pattern of staining, with a high density of immunolabeled *cilia* in the pyramidal stratum of CA1, CA2, and to a lesser extent, CA3 of the hippocampus proper. Scattered immunoreactivity was found in *strata Oriens* and *Radiatum.* The cortical distribution of MCHR1 followed a clear layer-specific distribution, with high densities of MCHR1 found in layers 2, 3 and 5, lower densities found in layers 4 and 6, and no staining found in layer 1 (Supplementary Material 01). Among the cortical regions containing MCHR1-ir *cilia* elements are: orbital, granular and agranular insular cortex; primary and secondary somatosensory cortices; primary and secondary motor cortices; frontal, perirhinal and entorhinal cortices; temporal, visual and auditory cortices. In the cingulate cortex, we observed a decreased density of MCHR1-ir *cilia* in layers 2 and 3 when compared to other cortical areas.

In the diencephalon, MCHR1-ir *cilia* were observed in several thalamic areas, such as the parataenial nucleus, the paraventricular thalamic nucleus, medial thalamic nuclei, and the medial habenular nucleus. We did not observe immunoreactivity in the *zona incerta.* In the hypothalamus, scattered *cilia* can be found in the lateral zone, in addition to strong clusters of immunoreactive material in the preoptic periventricular nucleus, paraventricular hypothalamic nucleus, and arcuate nucleus. Average densities of *cilia* were found in the medial preoptic area, anterior hypothalamic area and the posterior hypothalamic area, while medial zone nuclei were mostly devoid of labeling, such as the dorsomedial hypothalamic nucleus and the ventromedial hypothalamic nucleus.

At first glance, the distribution of MCHR1-ir *cilia* appeared to be complementary to that of MCH- and NEI-ir fibers, with areas rich in immunoreactive fibers virtually devoid of MCHR1-ir *cilia,* while areas rich in the receptor receive sparse innervation (Diniz *et al.* 2019). To confirm this pattern, we performed double immunohistochemistry to visualize MCHR1 and MCH-ir fibers in the same slices. The resulting pattern of labeling confirmed the segregation between some MCH-ir fibers-rich areas and areas containing large amounts of MCHR1-ir *cilia,* such as the nucleus accumbens/medial septal nucleus boundary (Fig. 5), or the medial/lateral *globus pallidus* (not shown).

**Figure 5 –.**
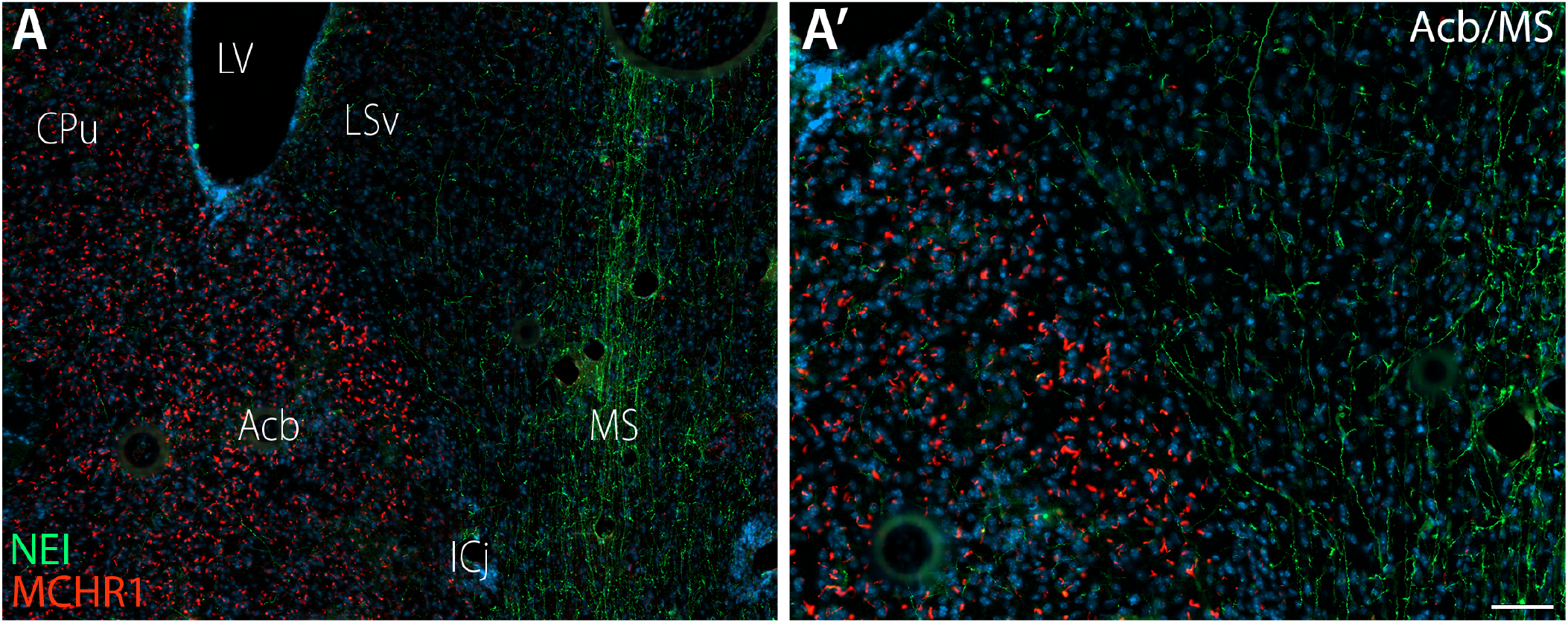
Segregation of MCH and MCHR1 fields. Widefield fluorescence photomicrographs of a frontal male mouse brain slice submitted to immunohistochemistry for MCH (green) and MCHR1 (red), and counterstained with DAPI (blue). A) The septal area shows a clear segregation between MCH-ir fibers and MCHR1-ir *cilia.* A high number of fibers occupy the medial septal nucleus and only sparely innervate the shell of the *nucleus accumbens,* while MCHR1-ir *cilia* are found in the *nucleus accumbens* but not in the medial septal nucleus; A’) Higher magnification of the delineated area in A. A clear MCH/MCHR1 boundary can be seen. Abbreviations: Acb – *nucleus accumbens*; CPu caudate-putamen; ICj – islands of Calleja; LSv – lateral septal nucleus, ventral part; LV – lateral ventral; MS – medial septal nucleus. Scale bar: A = 100μm; A’ = 50μm.

In rats, the general outline of staining was very similar to the one found in mice, including abundant staining in olfactory regions, chiefly the olfactory bulb and anterior olfactory nuclei, piriform cortex, olfactory tubercle, ventral *pallidum* and the islands of Calleja; the shell of the *nucleus accumbens;* layers 2, 3 and 5 of numerous cortical fields; the pyramidal stratum of the hippocampal formation; medial and central amygdaloid nuclei; midline thalamic nuclei and the periventricular and medial zones of the hypothalamus. Two notable differences, however, could be found between rats and mice. While mouse has a moderate-to-high staining of MCHR1 in the caudate putamen, rats were almost devoid of immunoreactivity is this area, with a very high background observed in the matrix. On the other hand, a negligible amount of MCHR1 immunoreactivity is observed in the mouse granular layer of the dentate gyrus, while a layer of stained *cilia* can be discerned in the subgranular zone of the rat dentate gyrus (Fig. 6).

**Figure 6 –.**
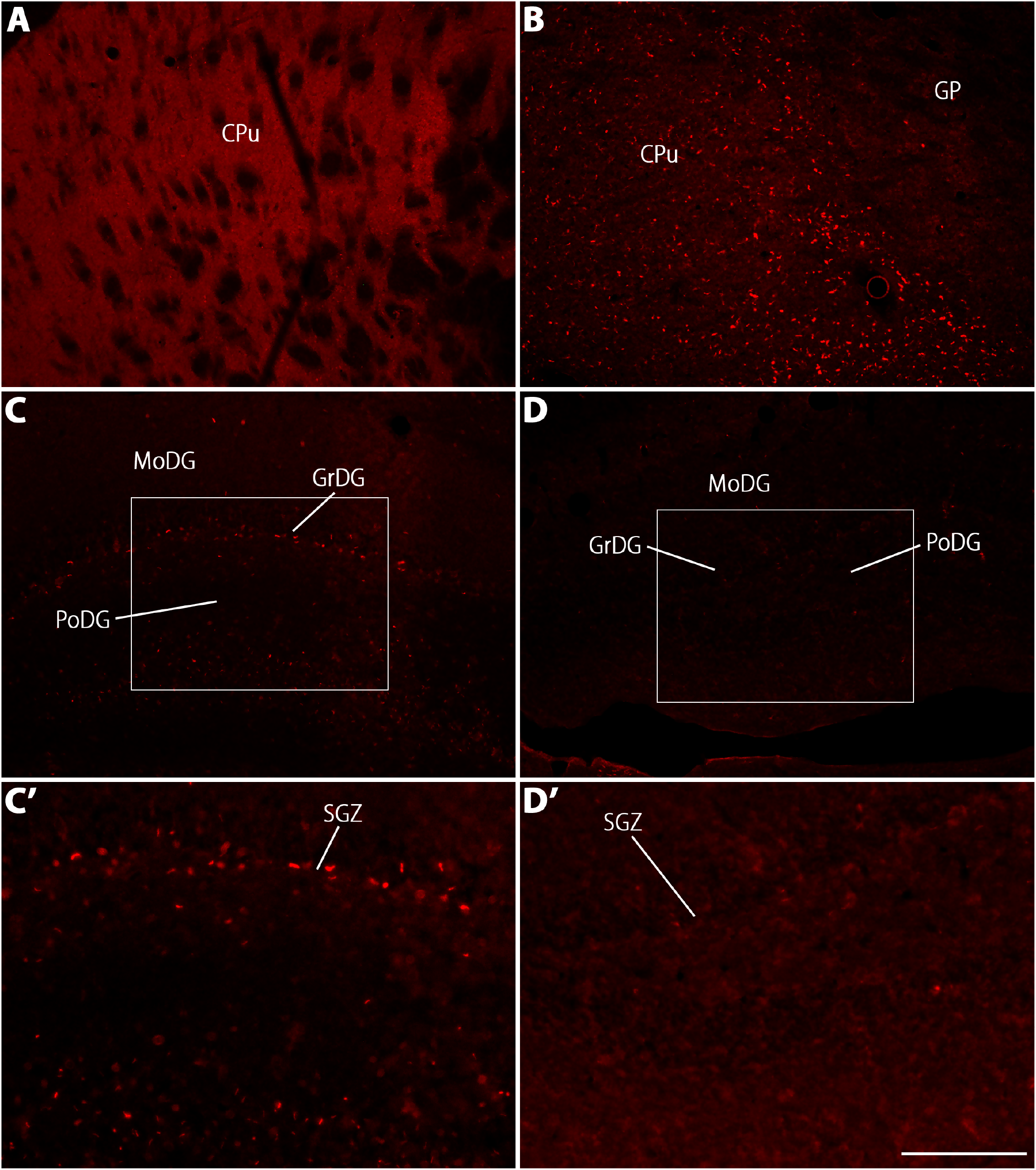
Differences between rats and mice concerning the distribution of MCHR1-ir *cilia.* Widefield fluorescence photomicrographs of frontal mice brain slices submitted to immunohistochemistry for MCHR1 (red). A, B) While the rat caudate-putamen displays a very high background and a very low number of immunoreactive *cilia,* the mouse caudate-putamen has a high number of immunoreactive *cilia.* C, D) The opposite is observed in the subgranular zone of the dentate gyrus. Although not abundant, rats display a strip of immunoreactive *cilia* in the subgranular zone, between the granular and polymorphic layers of the dentate gyrus. On the other hand, similar *cilia* are not observed in the mouse subgranular zone; C’, D’) Higher magnification of the delineated areas in C and D. Abbreviations: CPu – caudate-putamen; GP – *globus pallidus;* GrDG – granular layer of the dentate gyrus; MoDG – molecular layer of the dentate gyrus; PoDG – polymorphic layer of the dentate gyrus; SGZ – subgranular zone. Scale bar: A-D = 200μm; C’, D’ = 100μm.

### 3.3. Defined neurochemical populations display ciliary MCHR1

Because MCHR1-ir *cilia* were found in areas with heterogeneous neurochemical populations, we employed double immunohistochemistry to characterize some of those populations. In the olfactory bulb, MCHR1-ir *cilia* were found in a pattern that closely resembled the distribution of tyrosine hydroxylase-ir cells, bordering the glomeruli (Fig. 7 A, A’). To a lesser extent, MCHR1-ir *cilia* are found associated to calretinin-ir cells (Fig. 7 B, B’). On the other hand, no MCHR1-ir *cilia* were found within range of calbindin-ir cells (not shown). In the preoptic hypothalamus, we found MCHR1-ir *cilia* associated with kisspeptin-ir cells (Fig. 8 A, A’) but not with gonadotropin-releasing hormone (GnRH)-ir cells. In the periventricular and medial preoptic nuclei, MCHR1-ir *cilia* were also encountered in close proximity to estrogen receptor α cells (Fig. 8 B, B’).

**Figure 7 –.**
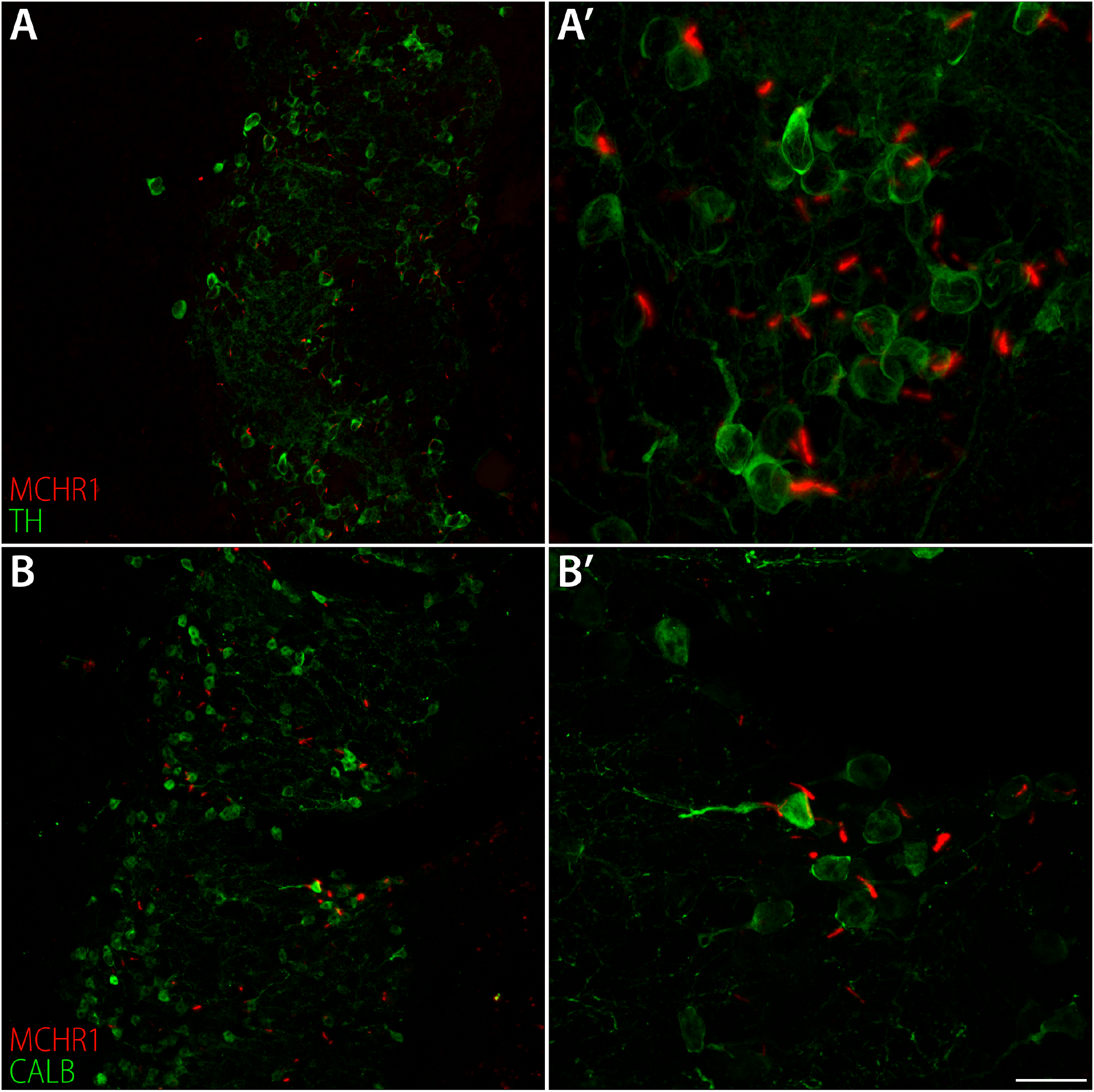
Neurochemistry identity of ciliary MCHR1-containing neurons in the glomerular layer of olfactory bulb. Confocal fluorescence photomicrographs of frontal mice brain slices submitted to immunohistochemistry for MCHR1 (red) and various neurochemical markers (green). A, A’) Ample colocalization between MCHR1 and tyrosine hydroxylase in the glomerular layer of the olfactory bulb; B, B’) Some colocalization is observed between calbindin and MCHR1, although in fewer numbers than MCHR1 and dopaminergic cells. Scale bar: A, B = 40μm; A’, B’ = 20μm.

In the paraventricular nucleus of the hypothalamus, MCHR1-ir *cilia* were densest in the parvicellular subdivisions, but immunoreactive *cilia* were also found in the magnocellular subdivisions. While there is no clear overlap on the MCHR1 and vasopressin distributions, MCHR1 is largely codistributed with oxytocin, mainly in the fringes of the magnocellular nuclei (Fig. 8 C, C’). A higher degree of colocalization was seen between MCHR1-ir *cilia* and corticotropin-releasing factor (CRF) in the parvicellular paraventricular nucleus (Fig. 8 D, D’). In the supraoptic nucleus, a different pattern was observed, with almost complete colocalization between MCHR1-ir *cilia* and vasopressinergic neurons, while only partial colocalization was found between MCHR1 and oxytocinergic neurons (Fig. 8 E, E’).

**Figure 8 –.**
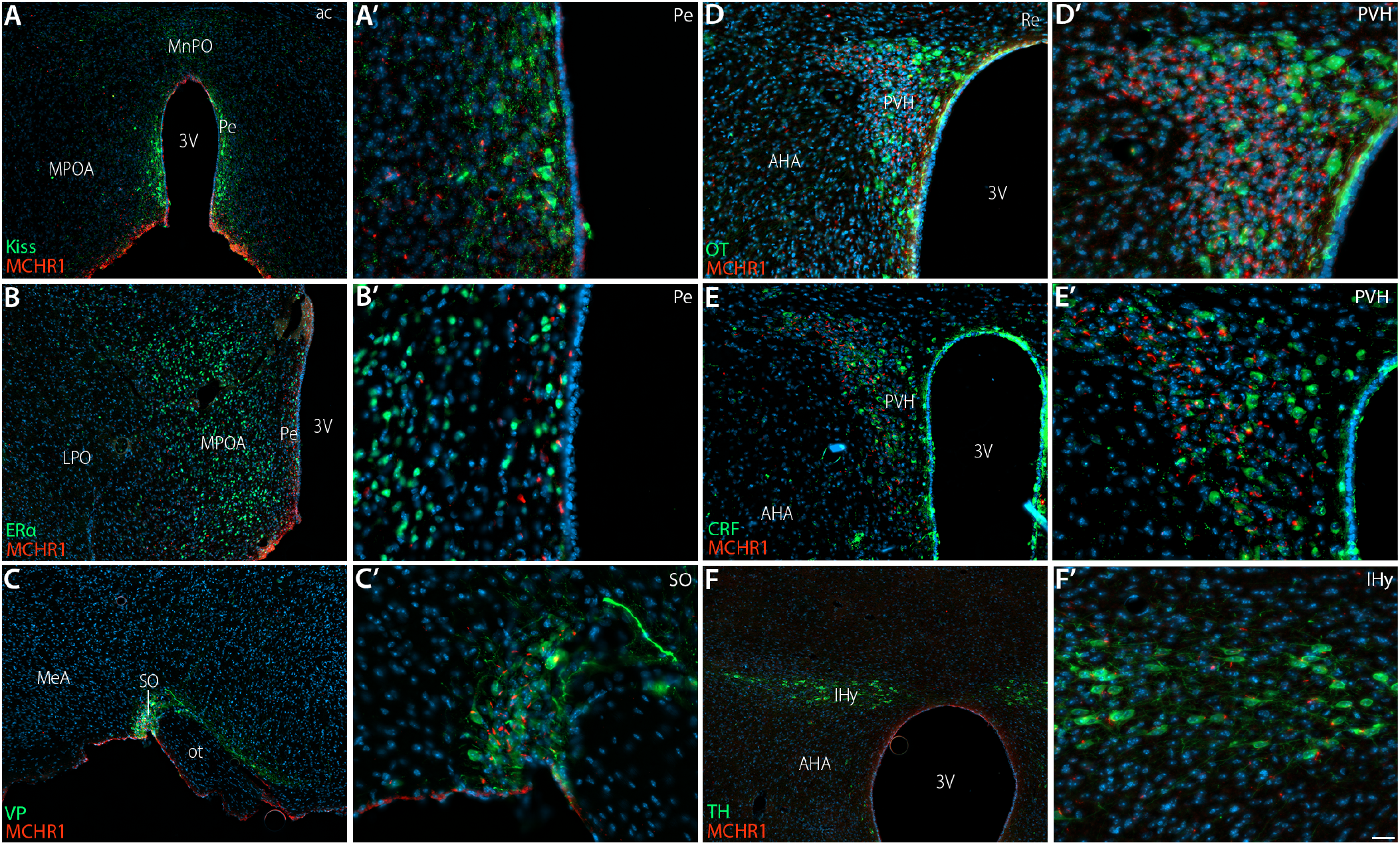
Neurochemistry identity of ciliary MCHR1-containing neurons in the hypothalamus. Widefield fluorescence photomicrographs of frontal mice brain slices submitted to immunohistochemistry for MCHR1 (red) and various neurochemical markers (green) counterstained with DAPI (blue). A, A’) In the periventricular nucleus, some kisspeptin-ir cells have MCHR1-ir *cilia;* B, B’) MCHR1-ir *cilia* are also seen in proximity to nuclei immunoreactive to estrogen receptor alpha in the periventricular nucleus; C – D’) In the paraventricular nucleus, MCHR1-ir *cilia* are found in oxytocin-ir and corticotropin-releasing factor-ir neurons; E, E’) In the supraoptic nucleus, the largest colocalization is between MCHR1-ir *cilia* and vasopressinergic cells; F, F’) In the incerto-hypothalamic area, virtually every dopaminergic cell has a MCHR1-ir *cilium.* Abbreviations: 3V – third ventricle; AHA – anterior hypothalamic area; IHy – incerto-hypothalamic area; LPO – lateral preoptic area; MeA – medial nucleus of the amygdala; MnPO – median preoptic nucleus; MPOA – medial preoptic nucleus; ot – optic tract; Pe – periventricular preoptic nucleus; PVH – paraventricular hypothalamic nucleus; Re – *nucleus reuniens*; SO-supraoptic nucleus. Scale bar: A-C, F = 100μm; D, E = 40μm; A’-F’ = 20μm.

In the arcuate nucleus, we found no colocalization between MCHR1 and α-melanocyte-stimulating hormone (α-MSH) or cocaine- and amphetamine-regulated transcript (CART). The only clear codistribution observed in the arcuate nucleus was between MCHR1 and tyrosine hydroxylase (TH), a dopaminergic marker. The arcuate nucleus, however, was not the only area where TH and MCHR1 were extensively codistributed, as virtually every TH-ir neuron in the incerto-hypothalamic area has a corresponding MCHR1-ir *cilium* (Fig. 8 F, F’). Neurons immunoreactive to CART in the incerto-hypothalamic area and in the lateral hypothalamus did not contain MCHR1-ir *cilia,* and there was no clear codistribution of MCHR1 and orexin in the lateral hypothalamus. Finally, no colocalization was observed between doublecortin and MCHR1 in the canonical sites of adult neurogenesis in the mouse brain.

## 4. Discussion

The presence of ciliary MCHR1 in the prosencephalon of murines is more extensive than previously known. In this work, we identified a commercial antibody that is able to stain ciliary MCHR1 with high specificity, and we employed it to map the presence of MCHR1 in neuronal primary *cilia* of both rats and mice, including female mice in all four stages of the reproductive cycle. Our results show a ciliary MCHR1 distribution that includes a wide range of structures, from the olfactory bulb to the hypothalamic mammillary nuclei. Our results are in agreement to most of what has been published up to this point in terms of *Mchr1* gene expression, suggesting ciliary MCHR1 is at least as abundant as, if not even more abundant, than other subcellular locations of MCHR1. The wide range of areas in which MCHR1 is found opens up the possibility that volume transmission may be a very important aspect of MCH cellular communication within the CNS, and an important aspect of neurochemical communication in vertebrate systems as a whole.

Up to this point, the staining of MCHR1 has been limited to a few works. Hervieu *et al.* (2000) were the first to use antibodies to map MCHR1, but there is no mention as to the subcellular localization of the labeling obtained by those authors. The other two published works that employed immunohistochemistry for MCHR1 were Berbari *et al.* (2008) and Niño-Rivero *et al.* (2019), who used a now discontinued antibody to demonstrate the presence of MCHR1 in the neuronal primary *cilia* of specific areas within the CNS of mice and rats. We demonstrated in this work that antibody PA5-24182 is a suitable replacement for the labeling of MCHR1, generating negative results in antibody omission tests, not labeling tissue from *Mchr1^-/-^*, and specifically labeling MCHR1 in neuronal primary *cilia,* as demonstrated by the colocalization with AC3. An additional benefit of the identified antibody is its target epitope. Directed at the last 27 residues of MCHR1, within the intracellular C-terminal portion of this receptor, this sequence is highly conserved between mammals, with a single conservative substitution in humans when compared to rodents, suggesting it may be successfully used to label MCHR1 in other mammals.

The distribution of MCHR1 obtained in this work is mostly similar, but not identical, to what has been described before. With respect to the immunohistochemical mapping of Hervieu *et al.* (2000) and the *in situ* hybridization mapping of Saito *et al.* (2001), several areas of the rat brain were also stained with antibody PA5-24182, including the olfactory tubercle, piriform cortex, endopiriform nuclei, hippocampal formation, amygdala, periventricular and paraventricular hypothalamic nuclei, arcuate nucleus, the cortical mantle and the midline thalamic nuclei. We also observed labeling in the olfactory bulb, islands of Calleja and in the medial habenular nucleus, areas that were also labeled in Hervieu *et al.* (2000), but not in Saito *et al.* (2001). On the other hand, Hervieu *et al.* (2000) described intense labeling of the medial septal nucleus and similar labeling of the hippocampus proper and the dentate gyrus, while this work and that of Saito *et al.* (2001) found an absence of MCHR1 synthesis and *Mchr1* expression in the medial septal nucleus, and a much lower presence of MCHR1/Mchr1 in the dentate gyrus when compared to the hippocampus proper.

The differences in MCHR1 density between rats and mice observed in this work are partially coherent with previous works. Hervieu *et al.* (2000) found a medium density of MCHR1 in the caudate-putamen of rats, while Saito *et al.* (2001) found a low expression of Mchr1 in this area. Chee *et al.* (2013) found a medium-high density of *Mchr1* reporter expression in mice. Our results align well with those of Chee *et al.* (2013) and Saito *et al.* (2001), as the mouse has a much more pronounced presence of MCHR1 *cilia* in the caudate-putamen. This is also the case with the dentate gyrus, as Saito *et al.* (2001) report a restricted presence of *Mchr1* expression in this structure, and Chee *et al.* (2013) found no *Mchr1* mRNA expression. To what extent these interspecies differences results in physiological differences is still unknown. Rats and mice have different profiles in terms of generation of new neurons in the hippocampus and their integration into functional circuits (Snyder *et al.* 2009), and the site of MCHR1 synthesis in rats found in this work is the subgranular zone, area thoroughly implicated in adult neurogenesis (Eriksson *et al.* 1998). Rats and mice also show different social responses to cocaine (Kummer *et al.* 2014), what could have as substrate differences in the caudate-putamen nucleus. To determine to what extent MCHR1 is involved in those differences more studies will be necessary.

Gene reporters and *in situ* hybridization have been used to investigate the expression of *Mchr1* in the mouse brain (Chee *et al.* 2013; Engle *et al.* 2018). Our distribution of MCHR1 immunoreactivity is mostly similar to what has been described in those works, including the olfactory areas, a similar pattern of cortical distribution, very dense labeling of the islands of Calleja, preeminent labeling in the pyramidal layer of the hippocampal formation and *induseum griseum,* no labeling of the medial septal nucleus, low labeling of thalamic nuclei, and weak to moderate labeling of the *zona incerta* and amygdala. The major disagreement between our work and that of Chee *et al.* (2013) concerns the relative density of synthesis/expression in some hypothalamic nuclei: while gene expression was highest in the arcuate nucleus and only moderate in the paraventricular and supraoptic nuclei, we found average synthesis in the arcuate nucleus and a very dense labeling density in the paraventricular and supraoptic nuclei. These disagreements likely stem from the inherent differences between the methods employed. Detection of *Mchr1* expression in the gene reporter model depends on the strength of *Cre* expression in the Mchr1-Cre lineage, which often is less than100%, resulting in neurons who are not detected despite expression of *Mchr1.* Furthermore, the density of Cre-positive neurons in gene reporter models cannot be correlated directly with protein synthesis, limiting a precise interpretation of MCHR1 function. Curiously, Engle *et al.* (2018) report that the level of *Mchr1* expression in the arcuate and paraventricular nuclei varied significantly between animals with inducible versus constitutive *Mchr1-Cre* alleles, suggesting that transient expression of *Mchr1* may occur during development at critical windows, possibly resulting in the variations observed between our animals and previous studies.

The labeling obtained in this work was entirely ciliary, as revealed by the full colocalization between MCHR1 and AC3 in all areas examined. It is unclear, at this moment, if this results from all MCHR1 being found solely at primary *cilia,* or if the antibody targets ciliary MCHR1 selectively, either due to slight post-translational differences between ciliary and non-ciliary MCHR1 or due to facilitated antibody access to its epitope in the *cilium* as compared to the membrane or synaptic space. Hervieu *et al.* (2000) found labeling with anti-MCHR1 to be confined to the plasma membrane, indicating that the antibody used in that work targets specifically membrane MCHR1, while both Berbari *et al.* (2008) and Niño-Rivero *et al.* (2019) found labeling to be exclusively ciliary. Regardless of the case, our work demonstrates that MCHR1 presence in the *cilium* is widespread, what has important functional implications. Neuronal primary *cilia* are believed to function as sensory organs for the cell (Pazour & Witman 2003). The existence of primary *cilia* as sensory organs can be explained through an evolutionary perspective: neuronal surface components tend to interfere with the homogenization of the ECS due to electrostatic interactions between the cellular membrane and ECS proteins. An appendicular structure that projects into the ECS, distancing itself from the cellular membrane, provides an adaptive advantage by better sensing subtle changes in the extracellular matrix composition (Marshall & Nonaka 2006). One direct inference that can be made from the fact that MCHR1 is found in the primary *cilia* is that MCH must be found in the ECS, outside the synaptic space, what is commonly associated with the concept of VT (Agnati *et al.* 1986; Agnati *et al.* 2010; Agnati *et al.* 1995).

Volume transmission concerns the communication between neurons that happens through pathways that are structurally poorly defined, occurs in a tridimensional space, and generally has multiple targets. Mechanisms of VT include the extrasynaptic spilling of neurotransmitters, release of neurochemical messengers in the ECS and CSF, and vesicle release, as opposed to wiring transmission, which includes synapses and gap junctions (Agnati & Fuxe 2014; Fuxe *et al.* 2007). Particularly relevant for ciliary receptors is the release of neurochemical messengers in the ECS and CSF. This release can occur in several ways, including peptide vesicle release at axonal non-synaptic domains (Golding 1994), reverse-uptake release of neurotransmitters (Attwell *et al.* 1993), and exocytotic and non-exocytotic somatodendritic release of peptides (Pow & Morris 1989). Once the signal is released, it can bind to receptors in the parent neurons, diffuse to neighbor neurons through the ECS, or travel to distant sites through the paravascular fluid circulation (Rennels *et al.* 1985), fiber bundle convection (Bjelke *et al.* 1995) or CSF flow.

Volume transmission mechanisms of communication have been demonstrated or suggested for a wide range of monoaminergic and peptidergic systems, including serotonin, norepinephrine, acetylcholine, dopamine, oxytocin, vasopressin, gonadotropin-releasing hormone, dynorphin, neurokinin A, substance P, cholecystokinin, enkephalin, and beta-endorphin, corticotropin-releasing factor, and orexin (Alpár *et al.* 2019; Agnati *et al.* 2010). Morphological aspects of those systems are often used as indicatives of VT, such as the presence of fiber plexuses close to the ventricle, fiber varicosities that lack synaptic specializations, somatodendritic release, and fiber-receptor mismatches. Recently, Noble *et al.* (2018) added MCH to the list of neuropeptides that use VT, by demonstrating that: MCH-ir fibers contact the ventricular space; 2. Levels of MCH in the CSF fluctuate tethered to MCH neuronal activation; 3. Activation of CSF-contacting fibers modulates feeding behavior. We recently demonstrated that the intimate relationship between MCH fibers and the ventricular space is a common feature of muroids, suggesting some degree of phylogenetic conservation, and that not only the ventricular *lumen,* but the subleptomeningeal space, may be used as a vehicle for the transport of MCH (Diniz *et al.* 2019). Given that MCH and NEI are found in different subcellular compartments, this opens up the possibility that the two peptides may be differentially released to act on wiring and VT paradigms (Diniz *et al.* 2019).

It was unclear, up to this point, what are the targets of the CSF-released MCH described by Noble *et al.* (2018). The results obtained in our work indicate that multiple regions are equipped to respond to free MCH, and that functions beyond the modulation of feeding behavior may be impacted. The two most likely targets for ingestive behavior modulation by free MCH are the arcuate nucleus and the paraventricular nucleus of the hypothalamus. Both structures have been thoroughly implicated in feeding behavior (Stanley & Leibowitz 1985; Bouret *et al.* 2004), and injections of MCH in these areas have been shown to modulate ingestion (Abbott *et al.* 2003). These two areas have some of the highest densities of ciliary MCHR1 in the murine diencephalon, making them particularly apt to detect free MCH, in particular the paraventricular nucleus due to its intimate association with the third ventricle. Although not as close the lateral ventricles as the paraventricular nucleus, numerous varicose MCH- and NEI-ir fibers are observed adjacent to the nucleus accumbens, suggesting MCH may be released by those fibers in the ECS and then diffuse to bind to ciliary MCHR1 in the *nucleus accumbens.* Although the arcuate nucleus would also be a possible target for ingestive modulation, we did not detect MCHR1 in the neurochemical populations of this area that have been related to feeding behavior.

Morphofunctional correlated indicate that several functions previously associated with MCH may be modulated through VT, in addition to the aforementioned role in feeding behavior. The widespread distribution of ciliary MCHR1 in the cortical mantle, including the motor and cingulate cortices, allows MCH to play a role in motor function (Segal-Lieberman *et al.* 2003) and the animal emotional state (Borowsky *et al.* 2002). The high density of MCHR1 in the caudate-putamen also allows MCH to act on reward circuits through the mesolimbic pathway (Domingos *et al.* 2013). Through the pyramidal layer of the hippocampus proper, MCH may act on learning, novelty, spatial and contextual memory retrieval, and anxiety (Sita *et al.* 2016; Monzon *et al.* 1999; Blanco-Centurion *et al.* 2019; Oh *et al.* 2019; Jimenez *et al.* 2018), with important ramifications for animal behavior. The presence of ciliary MCHR1 on dopaminergic neurons of the incerto-hypothalamic area may be the nexus for an MCH role on reproductive behavior and hormone secretion (Murray *et al.* 2000a), while through tuberoinfundibular dopaminergic neurons it may interface with prolactin secretion (Hökfelt & Fuxe 1972).

The arcuate nucleus is not the only pathway through which free MCH may modulate the adenohypophyseal release of hormones. Through the preoptic periventricular nucleus, free MCH from the CSF may act on kisspeptin neurons to modulate GnRH release (Wu *et al.* 2009), and in the PVH it may act to modulate the function of CRF neurons. While the former is likely related to a reproductive role (Murray *et al.* 2000b; Murray *et al.* 2000c; Murray *et al.* 2006), the modulation of CRF neurons may be related to control of stress responses and emotion (Lagos *et al.* 2011; Kennedy *et al.* 2003; Smith *et al.* 2006). A modulation of sexual function may also be achieved through estrogen receptor *α* neurons of the medial preoptic area and periventricular hypothalamus. The presence of MCHR1 in the medial preoptic area may also be important for the display of maternal behavior (Alachkar *et al.* 2016; Benedetto *et al.* 2014), if the lactation-generated MCH neurons of the medial preoptic area (Knollema *et al.* 1992; Rondini *et al.* 2010; Alvisi *et al.* 2016; Costa *et al.* 2019; Ferreira *et al.* 2017b) release MCH in a paracrine mode. Finally, another major function that may be played by MCH through VT is olfactory integration. One of the densest areas of ciliary MCHR1 is the olfactory bulb, where MCHR1 is found in several key layers, in addition to other ancillary areas important for olfactory processing, including the piriform cortex, anterior olfactory nucleus, olfactory tubercle, and *taenia tecta.* The presence of MCHR1 in two very important populations of the glomerular layer, dopamine and calbindin cells, in particular, indicates that those cells are the way through which MCH acts on olfactory integration, a role previously suggested for MCH (Adams *et al.* 2011; Alhassen *et al.* 2019).

In conclusion, we mapped for the first time the distribution of ciliary MCHR1 immunoreactivity in the brain of rats and mice. To do that, we evaluated a panel of commercial antibodies and identified one that has high specificity, good signal/background *ratio,* and that identifies an epitope that has been well conserved among mammals. The identification of immunoreactivity is a powerful tool, for it correlates with actual receptor production by the cells, giving us the closest functional readout of MCH action sites possible. All immunoreactivity detected with this antibody is ciliary, given the colocalization between MCHR1 and a specific neuronal primary *cilium* marker in all areas examined. Given the role of the neuronal primary *cilia* in sensing of the extracellular space, this colocalization suggests MCH can be found outside the synaptic space, and that free MCH may have an important role in VT modes of neuromodulatory signaling. Given the widespread distribution of ciliary MCHR1, it is fundamental that further studies are conducted to better understand how volume transmission affects the MCH system, what may open the possibility to new venues of pharmacological intervention targeting MCHR1 in the future.

**Supplementary Material 01 – Colocalization between MCHR1-ir *cilia* and NeuN in the cortex**. Widefield fluorescence photomicrographs of frontal mice brain slices submitted to immunohistochemistry for MCHR1 (red) and NeuN or DCX (green) counterstained with DAPI (blue). A, A’) *Cilia* immunoreactive to MCHR1 largely colocalizes with NeuN, reinforcing that primary *cilia*-containing cells are neurons. B, B’) There is no colocalization between MCHR1 and new neurons originated at the subgranular zone of the dentate gyrus. Abbreviations: 1 – cortical layer 1; 4 – cortical layer 4; 6 – cortical layer 6; cc – *corpus callosum*; GrDG – granular layer of the dentate gyrus; MoDG – molecular layer of the dentate gyrus; PoDG – polymorphic layer of the dentate gyrus. Scale bar: A = 100μm; B = 40μm; A’, B’ = 20μm.

