## Supplementary figures and images for "The distribution of neuronal primary *cilia* immunoreactive to melanin-concentrating hormone receptor 1 (MCHR1) in the murine prosencephalon"

### Supplementary Material 01

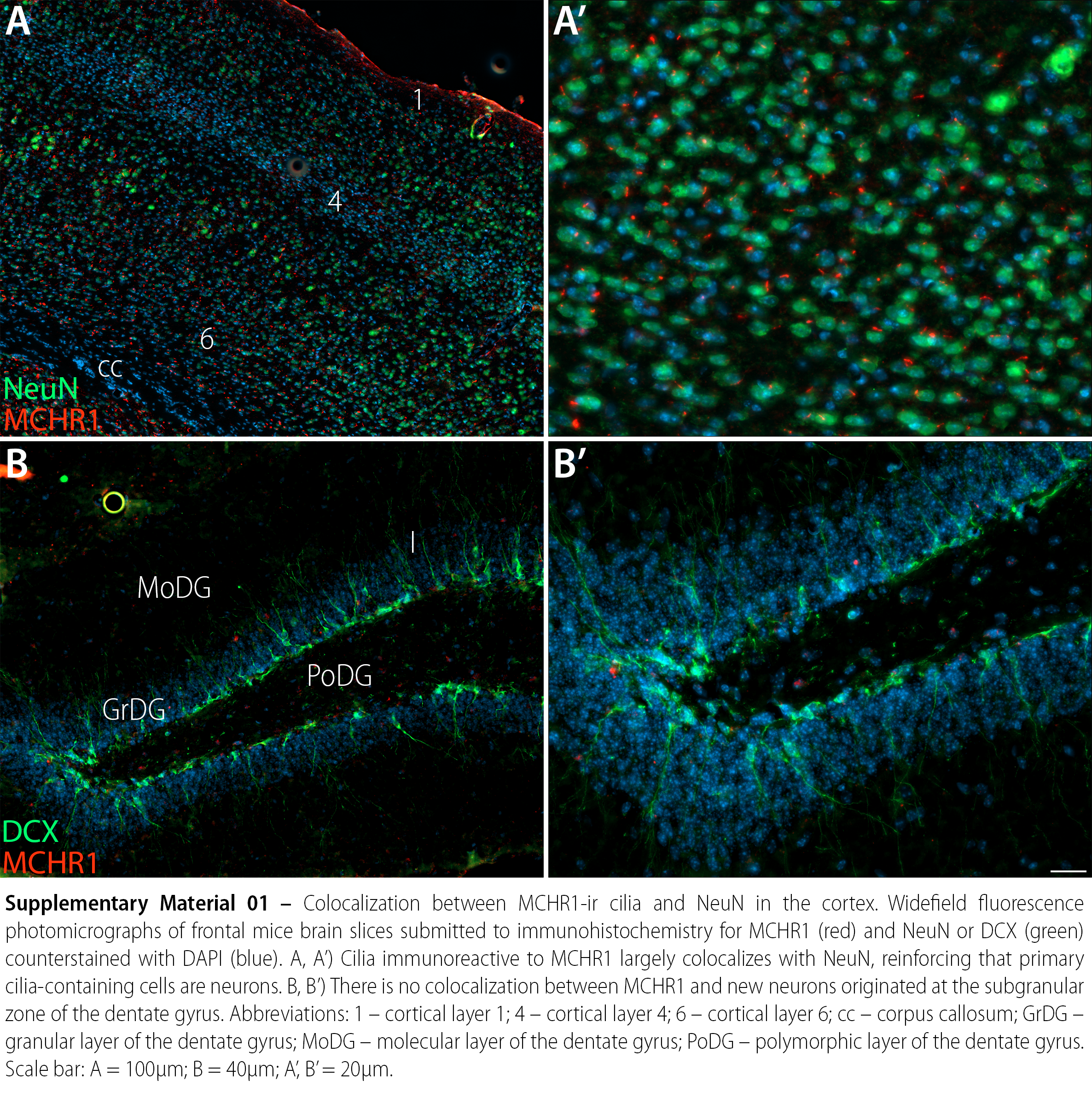
